# Bps polysaccharide of *Bordetella pertussis* resists antimicrobial peptides by functioning as a dual surface shield and decoy and converts *Escherichia coli* into a respiratory pathogen

**DOI:** 10.1101/2022.04.06.487353

**Authors:** Audra R. Fullen, Jessica L. Gutierrez-Ferman, Kacy S. Yount, Cheraton F. Love, Hyun G. Choi, Mario A. Vargas, Deepa Raju, Kara N. Corps, P. Lynne Howell, Purnima Dubey, Rajendar Deora

## Abstract

Infections and disease caused by the obligate human pathogen *Bordetella pertussis (Bp)* are increasing, despite widespread vaccinations. The current acellular pertussis vaccines remain ineffective against nasopharyngeal colonization, carriage, and transmission. In this work, we tested the hypothesis that Bordetella polysaccharide (Bps), a member of the poly-β-1,6-*N*-acetyl-D-glucosamine (PNAG/PGA) family of polysaccharides promotes respiratory tract colonization of *Bp* by resisting killing by antimicrobial peptides (AMPs). Genetic deletion of the *bpsA-D* locus, as well as treatment with the specific glycoside hydrolase Dispersin B, increased susceptibility to AMP-mediated killing. Bps was found to be both cell surface-associated and secreted during laboratory growth and mouse infections. Addition of bacterial supernatants containing Bps and purified Bps increased *B. pertussis* resistance to AMPs. By utilizing ELISA, immunoblot and flow cytometry assays, we show that Bps functions as a dual surface shield and decoy by inhibiting AMP binding. Co-inoculation of C57BL/6J mice with a Bps-proficient strain enhanced respiratory tract survival of the Bps-deficient strain. In combination, the presented results highlight the critical role of Bps as a central driver of *B. pertussis* pathogenesis. Heterologous production of Bps in a non-pathogenic *E. coli* K12 strain increased AMP resistance *in vitro,* and augmented bacterial survival and pathology in the mouse respiratory tract. Therefore, by conferring virulence traits across bacterial genera, Bps transforms a primarily intestinal and urinary tract bacterium into a respiratory pathogen. These studies can serve as a foundation for other PNAG/PGA polysaccharides and for the development of an effective *Bp* vaccine that includes Bps.

**Author summary:** Pertussis or whooping cough, caused by the obligate human pathogen *Bordetella pertussis (Bp),* is resurging in many countries. Currently, the mechanism by which *B. pertussis* subverts and resists host immunity is poorly known. In this manuscript, we examined the role of the *B. pertussis* polysaccharide Bps in promoting resistance to antimicrobial peptides (AMPs), a critical component of host immune defense. We show that the presence of Bps on the bacterial cell surface enhanced AMP resistance. Bps was secreted both during bacterial growth and during mouse infections. We further found that Bps functioned both as a surface shield and decoy, thereby inhibiting AMP binding. Simultaneous infection of mice with Bps-proficient and Bps- deficient strains resulted in greater survival of the Bps-deficient strain in the mouse respiratory tract. Finally, production of Bps in a non-pathogenic *E. coli* strain increased AMP resistance *in vitro,* and increased bacterial survival and heightened pathology in the mouse respiratory tract. Our study provides new insights into how *B. pertussis* has evolved to survive in the mammalian respiratory tract. Additionally, these studies underscore the potential of a single virulence factor to convert a non-pathogenic bacterium into a respiratory tract pathogen.

## Introduction

*Bordetella pertussis* (*Bp*) is a strict human-adapted Gram-negative respiratory tract pathogen, which causes whooping cough or pertussis, a highly contagious disease. Despite high and widespread vaccine coverage, the incidence of pertussis has increased in many countries. Pertussis is traditionally described as a childhood disease, which results in severe and sometimes fatal infections in newborns and infants. There has also been an increase in infections of vaccinated adolescents and adults, in whom pertussis manifests as a persistent cough with milder symptoms [1–4].

The adaptation of *Bp* to humans as an exclusive host was primarily associated with extensive loss of genomic content and gene inactivation [4, 5]. Thus, *Bp* is likely evolving to retain only genes required for its survival in human hosts. The *bpsA-D* locus, which encodes the *Bordetella* polysaccharide (Bps), is conserved in all sequenced and annotated strains of *Bp*. Therefore, investigating pathogenic roles of Bps will provide insights into how *Bp* survives in its only niche, the human respiratory tract. Bps belongs to the poly-β-1,6-*N*-acetyl-D-glucosamine (PNAG/PGA) family of polysaccharides. These polysaccharides are produced by numerous Gram-positive and Gram-negative bacteria, fungal and eukaryotic organisms including *Plasmodia* spp., the causative agents of malaria[6–9]. For *Bp*, Bps was the first identified factor to promote colonization of the mouse nose. In addition, Bps was also critical for the colonization of the mouse trachea and lungs [6]. However, the mechanism(s) by which Bps promotes respiratory tract survival is unclear.

After exposure to inhaled microorganisms, the innate immune system of the respiratory tract functions as a highly effective defense against infectious agents. Antimicrobial peptides (AMPs) are critical innate immune components within the respiratory tract which exhibit broad-spectrum and potent microbicidal activities against both Gram-negative and Gram-positive bacteria [10–12]. However, bacteria employ diverse strategies to resist killing by AMPs. Alterations of net charge and permeability of the cell surface are some commonly employed strategies for AMP resistance. Common bacterial factors in Gramnegative and Gram-positive bacteria that contribute to AMP resistance are LPS and teichoic acids (TA), respectively [13–15]. There are few reported mechanisms that detail how *Bp* resists human AMPs. Modification of the lipo-oligosaccharide lipid A region with glucosamine increases resistance against LL- 37 [16]. Similar to that observed for TA of Gram-positive bacteria and LPS of *V. cholerae* [15, 17–19], we described that the addition of the amino acid D-alanine to a yet unidentified outer membrane component enhances resistance to several human AMPs [20].

In the current manuscript, we tested the hypothesis that Bps promotes resistance of *Bp* to AMPs. We show that Bps binds AMPs and protects bacteria from AMP-mediated killing when present on the bacterial surface and as a secreted factor. We also investigated the pathogenic consequences of Bps production in a non-pathogenic commensal *E. coli* K12 strain. Production of Bps as the sole *Bp* factor was sufficient to confer on *E. coli* the ability to survive in the mouse respiratory tract and resulted in aggravated lung pathology.

## Results

### The *bpsA-D* polysaccharide locus of *B. pertussis* contributes to AMP resistance

We showed previously that a *Bp* strain harboring an in-frame deletion in the genes of the *bpsA-D* locus (Δ*bpsA-D*), which encodes the Bps polysaccharide, colonized the mouse respiratory tract less efficiently than the wildtype (WT) strain as early as six hours post-challenge [6]. Since AMPs act early after infection to protect the respiratory tract from bacterial pathogens, we hypothesized that the Δ*bpsA-D* strain will be more sensitive to these antibacterial compounds than the WT strain. We first compared the sensitivities of these two strains to polymyxin B (PmB), a cationic antibiotic peptide that is an excellent model for bactericidal actions of AMPs [21]. Compared to the WT strain, the *ΔbpsA-D* mutant was significantly more sensitive to PmB-mediated killing (Fig. 1A). Complementation of the Δ*bpsA-D* mutant with the plasmid pMM11 containing the cloned *bpsA-D* locus (Δ*bpsA-D*^comp^) considerably increased bacterial survival in the presence of PmB when compared to that of the mutant strain harboring the vector plasmid only (Δ*bpsA-D*^vec^) (Fig. 1A).

**Figure 1.**
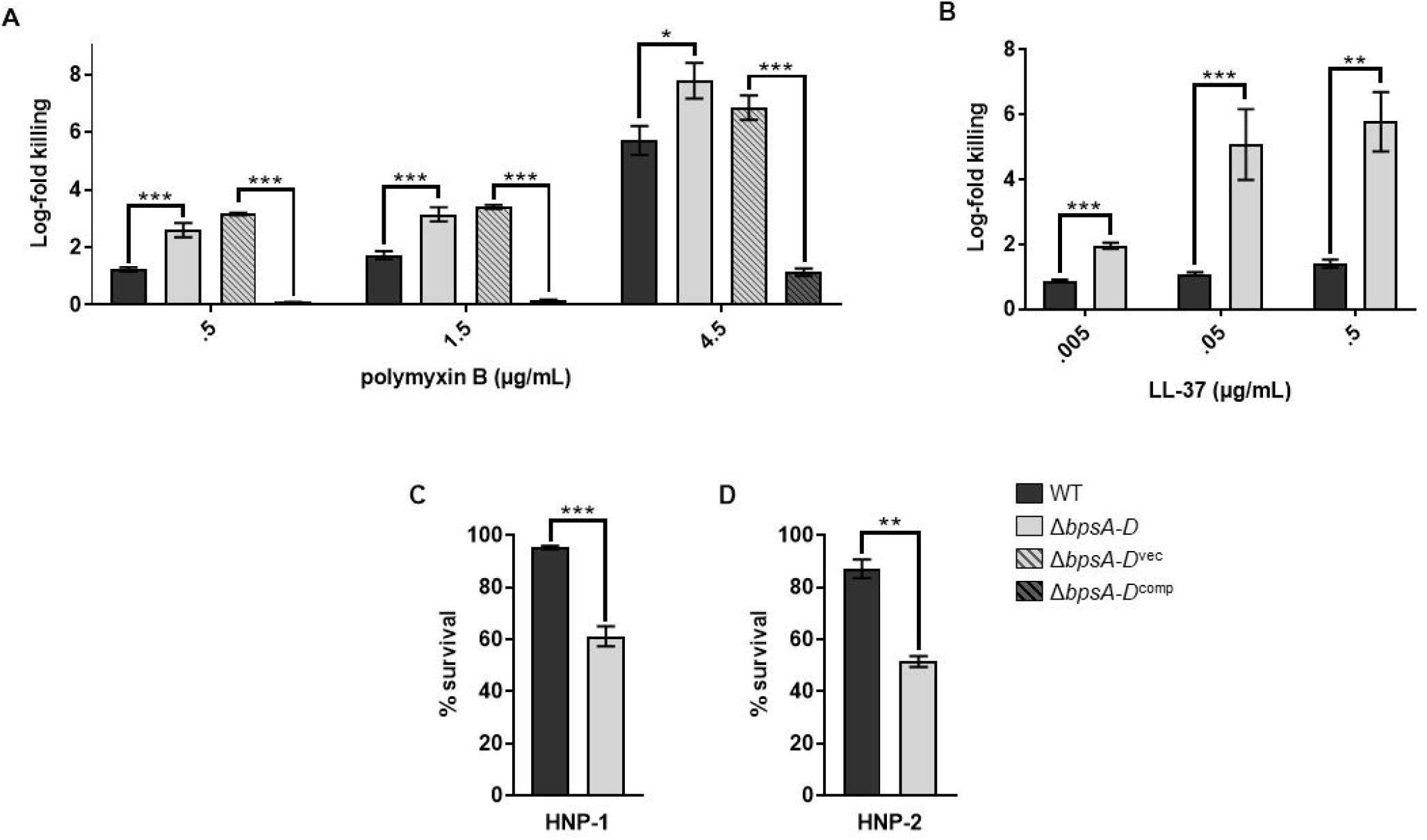
Bps promotes resistance to polymyxin B and LL-37. Survival of WT, *ΔbpsA-D, ΔbpsA-D^vec^*, and Δ*bpsA-D*^comp^ strains in the presence of polymyxin B (a), LL-37 (b), HNP-1 (c), and HNP-2 (d). Bacteria were exposed to the indicated concentrations of AMPs in 10mM Na3PO4 for 2 hours. Each data point represents the mean and s.e.m. of triplicates from one experiment and is representative of at least three independent experiments. Statistical differences were assessed by unpaired two-tailed Student’s *t* test. *, p<0.05; **, p<0.005; ***, p<0.0005.

Next, we tested the sensitivities of the WT and *ΔbpsA-D* strains to human AMPs. We used LL-37 (a human cathelicidin produced by phagocytes and epithelial cells in the respiratory tract) [22] along with HNP-1 and HNP-2 (members of α-defensin family found in the human respiratory tract) [23, 24]. Compared to the WT strain, the *ΔbpsA-D* mutant was killed significantly better in the presence of all these host AMPs (Fig. 1B-D). Taken together, these results demonstrate that the *bpsA-D* locus promotes resistance to several different AMPs.

### Dispersin B increases the susceptibility of *B. pertussis* to PmB and LL-37

Dispersin B (DspB) is a glycoside hydrolase that specifically hydrolyzes poly-β-1,6-N-acetyl-D-glucosamines [25–27]. While the exact structure of Bps is unknown, based on its immune reactivity and susceptibility to DspB, Bps appears to be a poly-β-1-6-N-acetyl-D-glucosamine polysaccharide [6, 27, 28]. Thus, DspB offers a useful biochemical tool to assess the contribution of Bps to AMP resistance independent of the *ΔbpsA-D* strain.

The WT and *ΔbpsA-D* strains were treated with various concentrations of DspB before incubation with PmB or LL-37. Based on the results in Fig. 2A, 50 μg/ml of DspB was chosen for further experiments, since it reduced the levels of detectable Bps on WT bacteria to that detectable on *ΔbpsA-D* bacteria. Compared to incubation with PmB or LL-37 alone, treatment of the WT strain with either DspB + PmB (Fig. 2B) or DspB + LL-37 (Fig. 2C) resulted in a 3.5- and 3.0-log-fold increase in bacterial killing, respectively. Compared to treatment with PmB or LL-37 alone, treatment of the *ΔbpsA-D* strain with DspB + PmB (Fig. 2B) or DspB + LL-37 (Fig. 2C) did not result in any further increase in bacterial killing, suggesting that the activity of DspB is specific to Bps. Treatment with DspB alone did not have any significant effect on the survival of either the WT or *ΔbpsA-D* strains (Fig. S1), suggesting that DspB does not have any toxic effect on *Bp*. These data show that enzymatic degradation of Bps increases the susceptibility of *Bp* to killing by PmB and LL-37.

**Figure 2.**
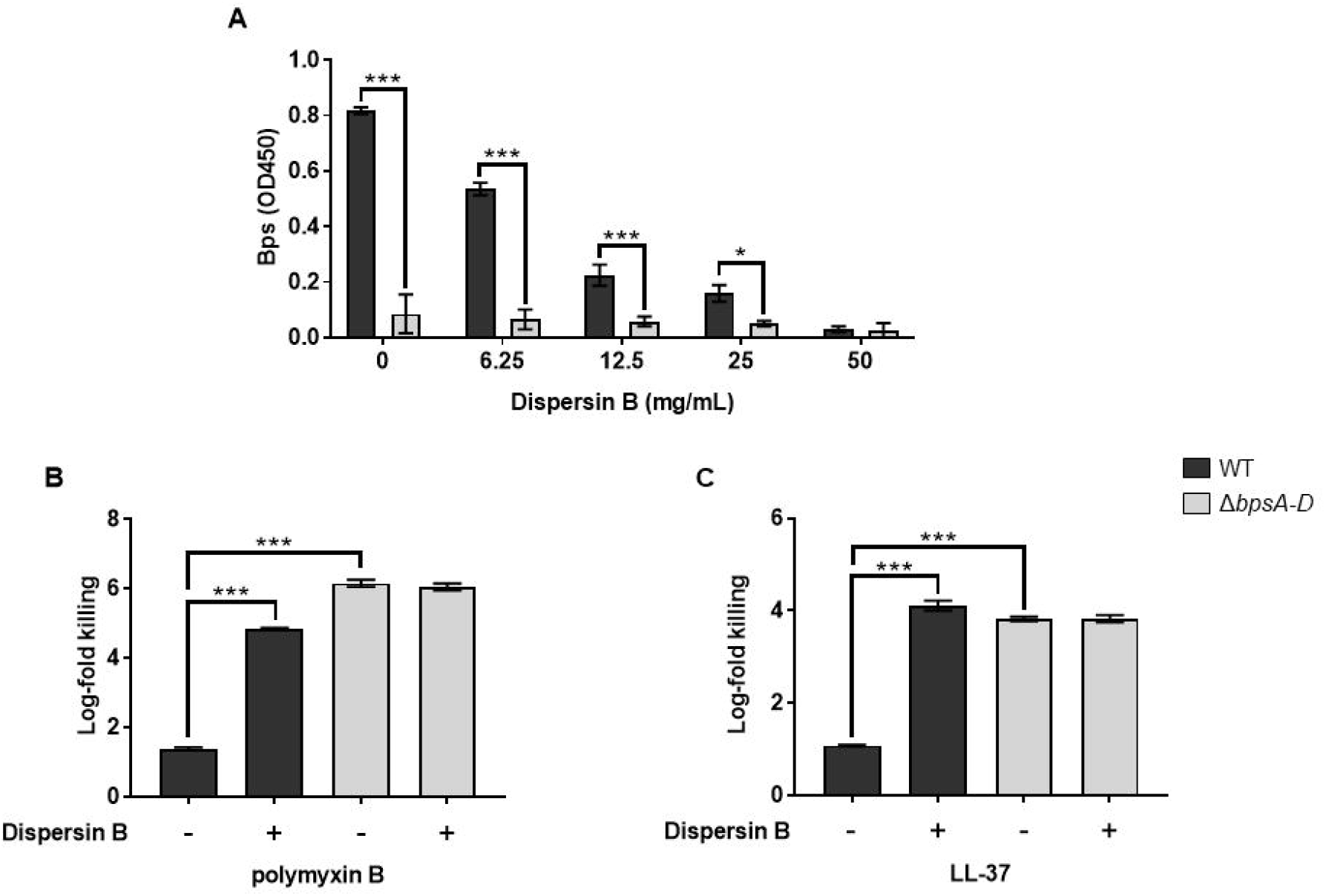
Dispersin B hydrolyzes Bps and increases the susceptibility of *B. pertussis* to PmB and LL- 37. (a) Bps was quantified by ELISA from the WT or *ΔbpsA-D* strains following incubation with indicated concentrations of Dispersin B. (b, c) Survival of WT and *ΔbpsA-D* strains following treatment with 50 μg/ml Dispersin B or buffer in the presence of 0.5 μg/ml polymyxin B (b) or 0.5 μg/ml LL-37 (c). Each data point represents the mean and s.e.m. of triplicates from one experiment and is representative of two independent experiments. Statistical differences were assessed by two-way ANOVA. *, p<0.05; ***, p<0.0005.

### The presence of Bps on the *B. pertussis* cell surface inhibits LL-37 binding

Since the results presented thus far suggest that Bps protects *Bp* by limiting the killing activity of PmB and LL-37, we hypothesized that Bps inhibits the binding of AMP to the bacterial cell surface. To test this, we used flow cytometry to quantify the binding of FITC-labelled LL-37 to fixed bacteria. Fig. 3A shows overlaid histograms comparing median fluorescence intensities of FITC, which are quantified in Fig. 3B. Addition of FITC- labeled LL-37 to WT bacteria resulted in an increase in FITC fluorescence compared to control (WT bacteria alone), indicating that LL-37 binds to WT bacteria. Addition of FITC-labeled LL-37 to *ΔbpsA-D* bacteria led to a considerable increase in FITC fluorescence, suggesting that higher amounts of LL-37 bind to the *ΔbpsA-D* strain than to the WT strain. These results suggest that the presence of Bps on the *Bp* cell surface reduces the binding of LL-37, providing one explanation for reduced AMP-mediated killing of the WT bacteria.

**Figure 3.**
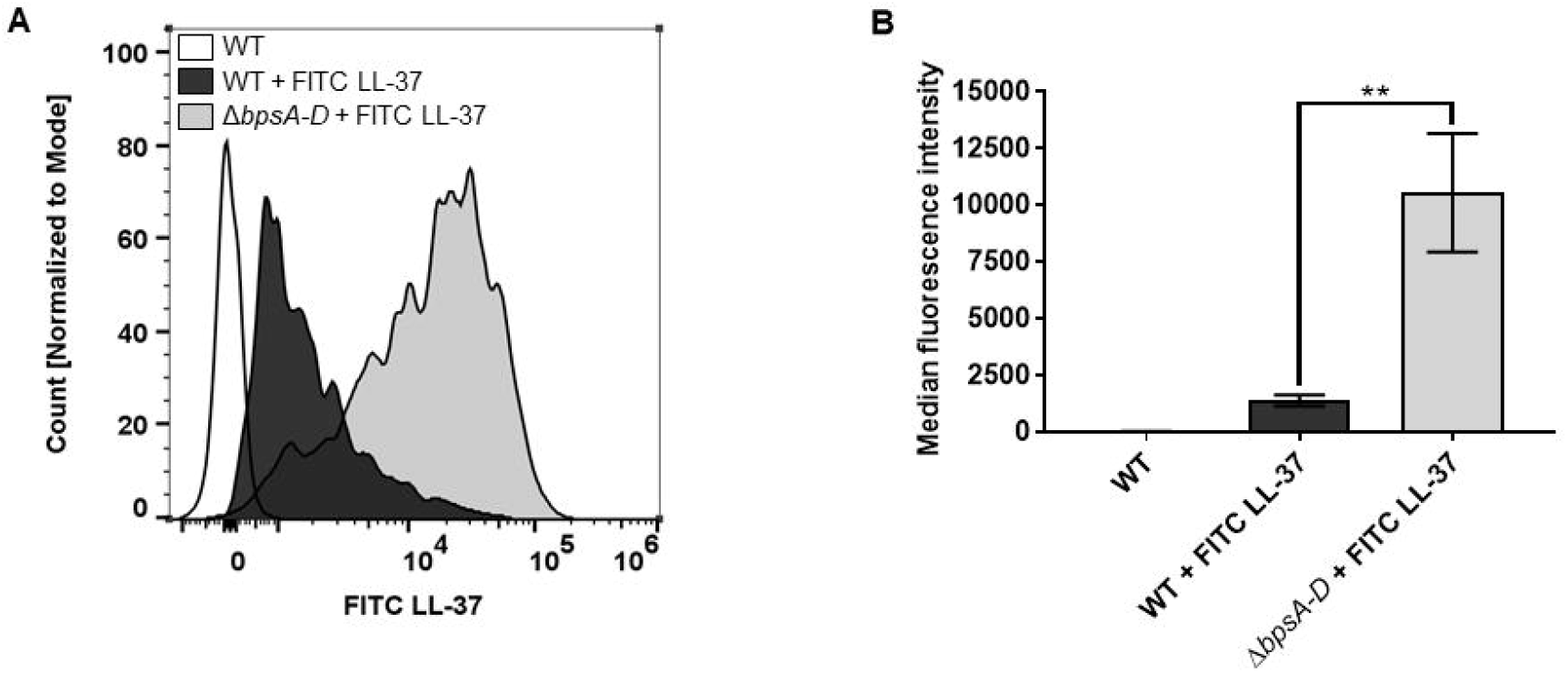
The presence of Bps on the cell surface inhibits LL-37 binding. (a, b) Bacteria were fixed and exposed to FITC-labeled LL-37 followed by measurement of the fluorescence intensity of FITC-labeled LL-37 bound to bacteria by flow cytometry. (a) Bacterial counts for each strain were normalized to mode. (b) Median fluorescence intensities were quantified and assessed for statistical analysis. Data represent triplicates from one of two independent experiments. Statistical differences were assessed by one-way ANOVA. **, p<0.005.

### Bps is secreted during laboratory growth of *B. pertussis*

In addition to producing polysaccharides on the cell surface, many bacteria also secrete them in the growth medium [8, 29]. It is not known if Bps is naturally secreted during laboratory growth of *Bp*. To determine this, the amounts of Bps present on the bacterial cell surface (cell-associated) and released into the growth medium (supernatant; collected after centrifugation and filtration of the spent medium) were quantitated by ELISA using the lectin wheat germ agglutinin (WGA) conjugated to HRP. In WT cells, Bps was detected on both the cell surface and in the supernatant (Fig. 4A). As expected, negligible amounts of Bps were detected on the cell-surface or in supernatant obtained from the *ΔbpsA-D* strain. These results demonstrate that *Bp* secretes Bps during laboratory growth.

**Figure 4.**
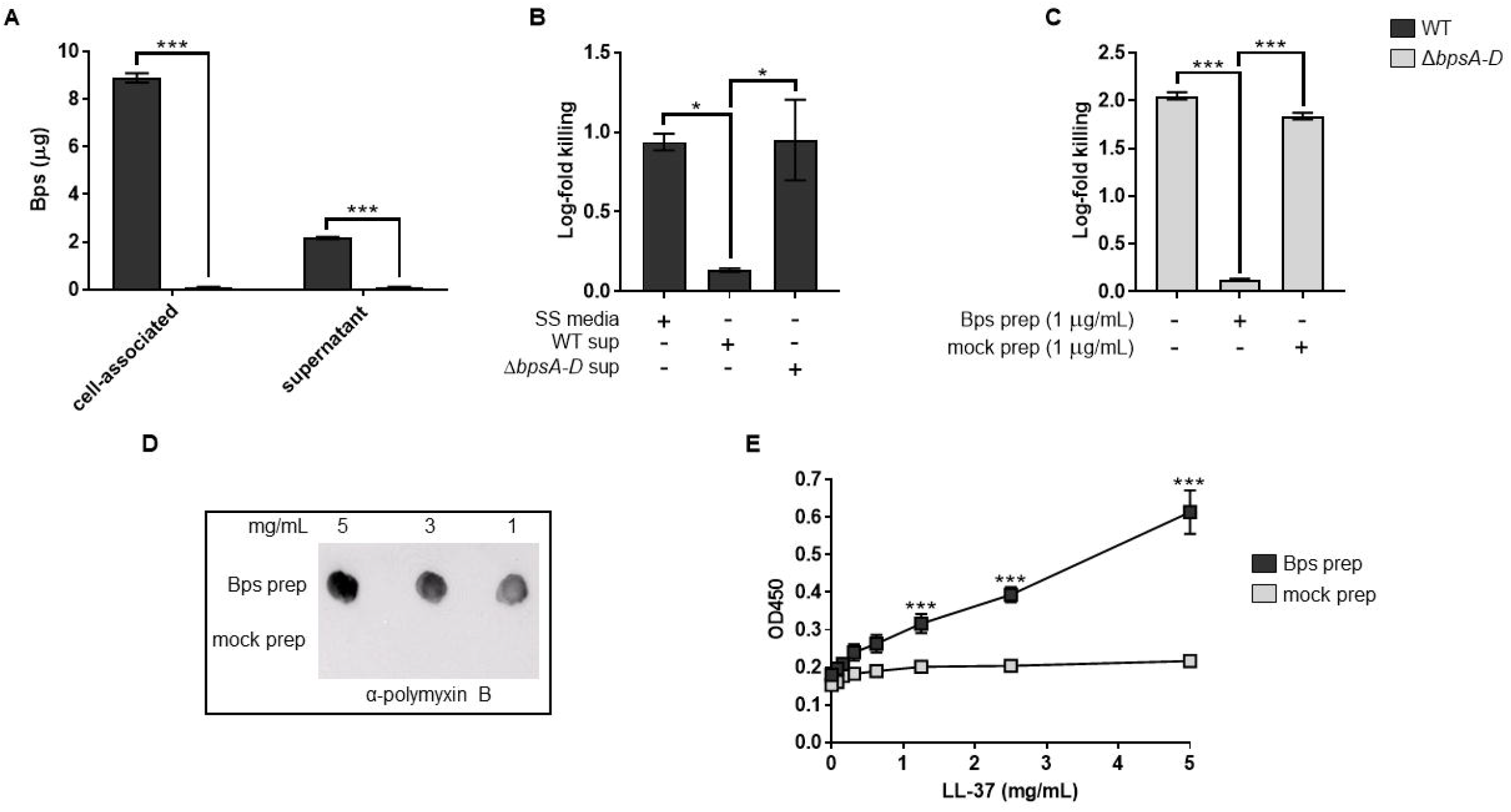
*B. pertussis* secretes Bps during laboratory growth, and cell-free Bps provides protection from and binds to PmB and LL-37. (a) Quantitation of cell-associated and secreted (supernatant) Bps from the WT and *ΔbpsA-D* strains by ELISA. Each data point represents the mean and s.e.m. of eight wells from one experiment and is representative of two independent experiments. Statistical differences were assessed by two-way ANOVA. ***, p<0.0005. (b, c) Survival of WT and *ΔbpsA-D* strains in the presence of PmB (b) or LL-37 (c). Supernatants from WT and the Δ*bpsA-D* cultures (b) or purified Bps and mock preparation (c) were added as indicated. Each data point represents the mean and s.e.m. of triplicates from one experiment and is representative of two independent experiments. Statistical differences were assessed by one-way ANOVA. *, p<0.05 ***, p<0.0005. (d, e) Bps polysaccharide binds PmB and LL-37. (d) Bps or mock preparations were spotted on nitrocellulose membranes. To detect PmB binding, the membranes were incubated with PmB, washed and probed with α-polymyxin B antibody conjugated to HRP. (e) Binding of Bps or mock preparations to LL- 37 was quantified by ELISA using WGA conjugated to HRP. Asterisks indicate significance compared to mock prep. Each data point represents the mean and s.e.m. from one experiment and is representative of two independent experiments. Statistical differences between Bps and mock preps were assessed by unpaired two-tailed Student’s *t* test. ***, p<0.0005.

### Cell-free Bps increases the resistance of *B. pertussis* to AMPs

Next, we determined if Bps secreted into the culture medium would increase AMP resistance. First, WT bacteria were incubated with PmB in the presence of SS media or filtered supernatants from either the WT or *ΔbpsA-D* bacteria. Compared to the addition of either SS media or supernatant from the *ΔbpsA-D* strain *(ΔbpsA-D* sup), addition of supernatant from the WT strain (WT sup) resulted in lower killing of the WT strain by PmB (Fig. 4B). We also tested if Bps purified from bacterial cells would increase bacterial survival. Addition of purified Bps (Bps prep) decreased killing of the *ΔbpsA-D* bacteria in the presence of LL-37. In contrast, addition of a mock-purified preparation from the *ΔbpsA-D* strain did not have any significant effect on the survival of the *ΔbpsA-D* strain (Fig. 4C). Collectively, these results suggest that the addition of secreted Bps increases AMP resistance.

### Cell-free Bps binds to PmB and LL-37

We used immunoblotting and ELISA to test whether cell-free Bps bound PmB and LL-37, respectively. For immunoblots, different amounts of Bps prep or mock prep were spotted on a nitrocellulose membrane, followed by incubation with PmB. After extensive washing, the bound PmB was detected by incubation with α-PmB antibody followed by an appropriate HRP- conjugated secondary antibody. PmB bound to the Bps prep but not to the mock prep (Fig. 4D). Similarly, LL-37 bound to Bps in a dose-dependent manner as detected by ELISA, whereas the mock prep showed very weak binding which did not increase with increasing amounts of LL-37 (Fig. 4E). Collectively, these results suggest that cell-free Bps limits the killing activity of AMPs by binding to them.

### Presence of the WT strain increases resistance to LL-37-mediated killing *in vitro* and promotes respiratory tract survival of the *ΔbpsA-D* strain

Based on the finding that cell-free Bps protects against AMP-mediated killing, we hypothesized that the WT strain will protect the *ΔbpsA-D* strain from AMP- mediated killing. To test this *in vitro,* we incubated monocultures or co-cultures (1:1) of the two strains with LL-37 and enumerated CFUs. When incubated with the WT strain, the susceptibility of the *ΔbpsA-D* strain to LL-37 was nearly identical to that of the WT strain (Fig. 5A).

**Figure 5.**
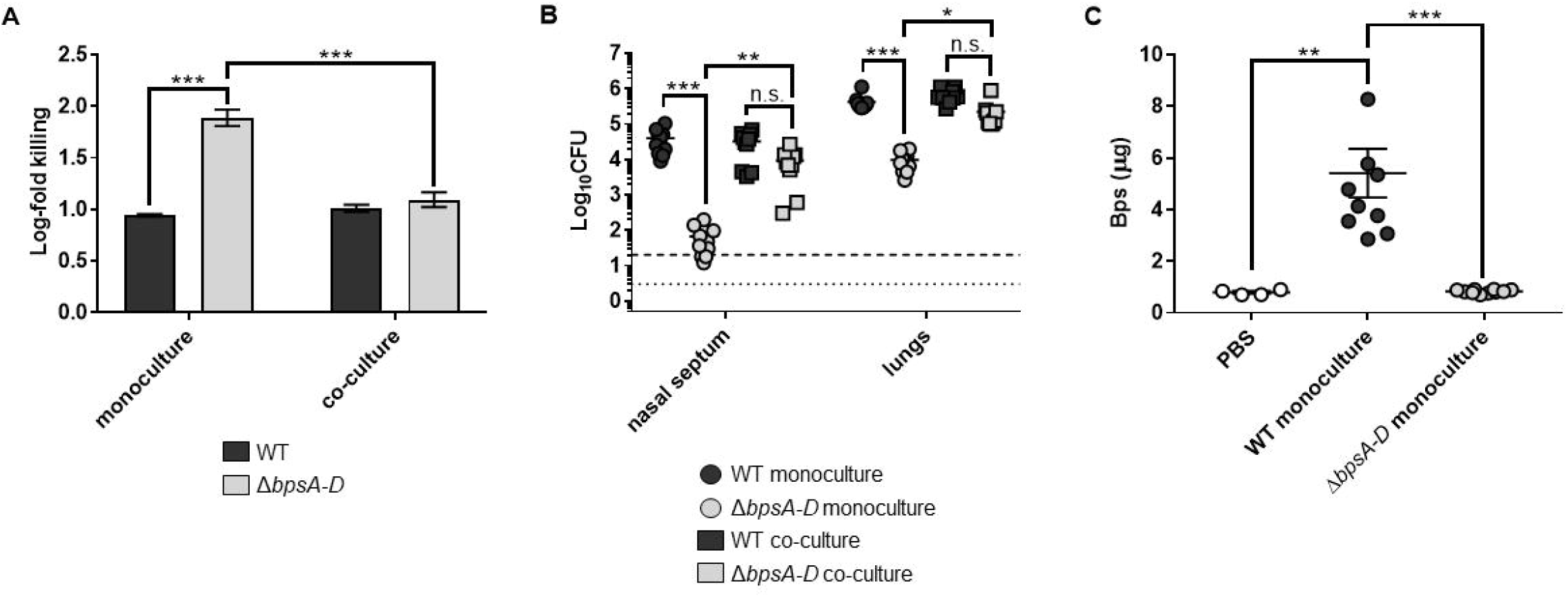
The presence of the WT strain increases *ΔbpsA-D* resistance to LL-37 *in vitro,* and enhances its survival in the mouse respiratory tract and Bps is secreted in mouse lungs during infection. (a) Survival of WT and *ΔbpsA-D* strains either in monoculture or in a 1:1 co-culture of both strains against LL- 37. Each data point represents the mean and s.e.m. of triplicates from one of five independent experiments. Statistical differences were assessed by two-way ANOVA. ***, p<0.0005. (b) Bacterial CFUs recovered from the nasal septum and lungs of C57BL/6J mice four days after aerosol infection with WT or *ΔbpsA-D* in monoculture or in a 1:1 co-culture. Bars indicate the mean and s.e.m. of ten mice. Data from two independent experiments with groups of five mice each are shown. Statistical differences were assessed by two-way ANOVA. n.s., not significant; *, p<0.05; **, p<0.005; ***, p<0.0005. Dotted line represents the lower limit of detection for nasal septum, and dashed line represents the lower limit of detection for lungs. (c) Amounts of Bps in supernatants of lung lysates obtained from mice either instilled with PBS (blank circles) or infected with the indicated strains was quantified by ELISA using WGA conjugated to HRP. For infected mice, bars indicate the mean and s.e.m. of two independent experiments consisting of groups of five mice each (from Fig. 5B). For PBS-instilled mice, bars indicate the mean and s.e.m. of one experiment consisting of four mice. Statistical differences were assessed by one-way ANOVA. **, p<0.005; ***, p<0.0005.

We then tested if a similar survival advantage will exist for the *ΔbpsA-D* mutant in the mouse respiratory tract when co-infected with the WT strain. C57BL/6J mice were infected by aerosol exposure with the WT or the *ΔbpsA-D* strain either as single strains or when combined in a 1:1 ratio. Bacterial burden was determined by enumeration of CFUs from the nasal septum and lungs at four days post-challenge. Enumeration of CFUs approximately 30 minutes after aerosol challenge confirmed that similar numbers of the two strains were delivered into the nose and the lungs (Fig. S2). Mice infected with only the *ΔbpsA-D* strain (gray circles) harbored significantly lower bacterial burden on the nasal septum and in the lungs compared to mice infected with the WT strain alone (black circles) (Fig. 5B). This result is consistent with our previously published results using the intranasal challenge route [6]. Strikingly, when co-infected with the WT strain, the bacterial burden of the Δ*bpsA-D* strain (grey squares) on the nasal septum and in the lungs was similar to that of the WT strain (black squares; Fig. 5B). These results indicate that the presence of the WT strain supports the colonization of the Δ*bpsA-D* strain in the mouse nose and lungs.

### *B. pertussis* secretes Bps in the mouse lungs

Lung lysates from mice were centrifuged and filtered, followed by quantification of Bps in the supernatants by ELISA. Bps was detected in lung supernatants of mice infected with the WT strain (black circles; Fig. 5C). In comparison, Bps levels in the lung lysates of mice infected with the *ΔbpsA-D* strain (gray circles) were similar to those of mice instilled only with PBS (clear circles). We speculate that this background level of Bps reactivity is due to the cross-reactivity of the WGA with host carbohydrates [30]. These results suggest that Bps secreted from the WT strain contributes to the increased bacterial burden of the *ΔbpsA-D* strain in the mouse respiratory tract.

### Production of Bps in *E. coli* confers resistance to PmB and LL-37, enhances bacterial survival in the mouse respiratory tract, and augments pathology in the lungs

We then tested whether Bps as the sole *Bordetella* factor was sufficient to provide resistance to AMP-mediated killing in *E. coli*. The pMM11 plasmid which encodes Bps was transformed into a derivative of MG1655 (*E. coli* K12 strain) [31], denoted as ARF001. Compared to the strain containing the empty vector (ARF001^vec^), the strain containing pMM11 (ARF001^*bpsA-D*^) was killed at lower numbers in the presence of PmB (Fig. 6A) and LL-37 (Fig. 6B). We also confirmed using FITC-labeled LL-37 that production of Bps in *E. coli* inhibited AMP binding (Fig. 6C-D). These results suggest that Bps confers resistance to PmB and LL-37 in *E. coli* independent of other *Bp* factors by inhibiting their binding.

**Figure 6.**
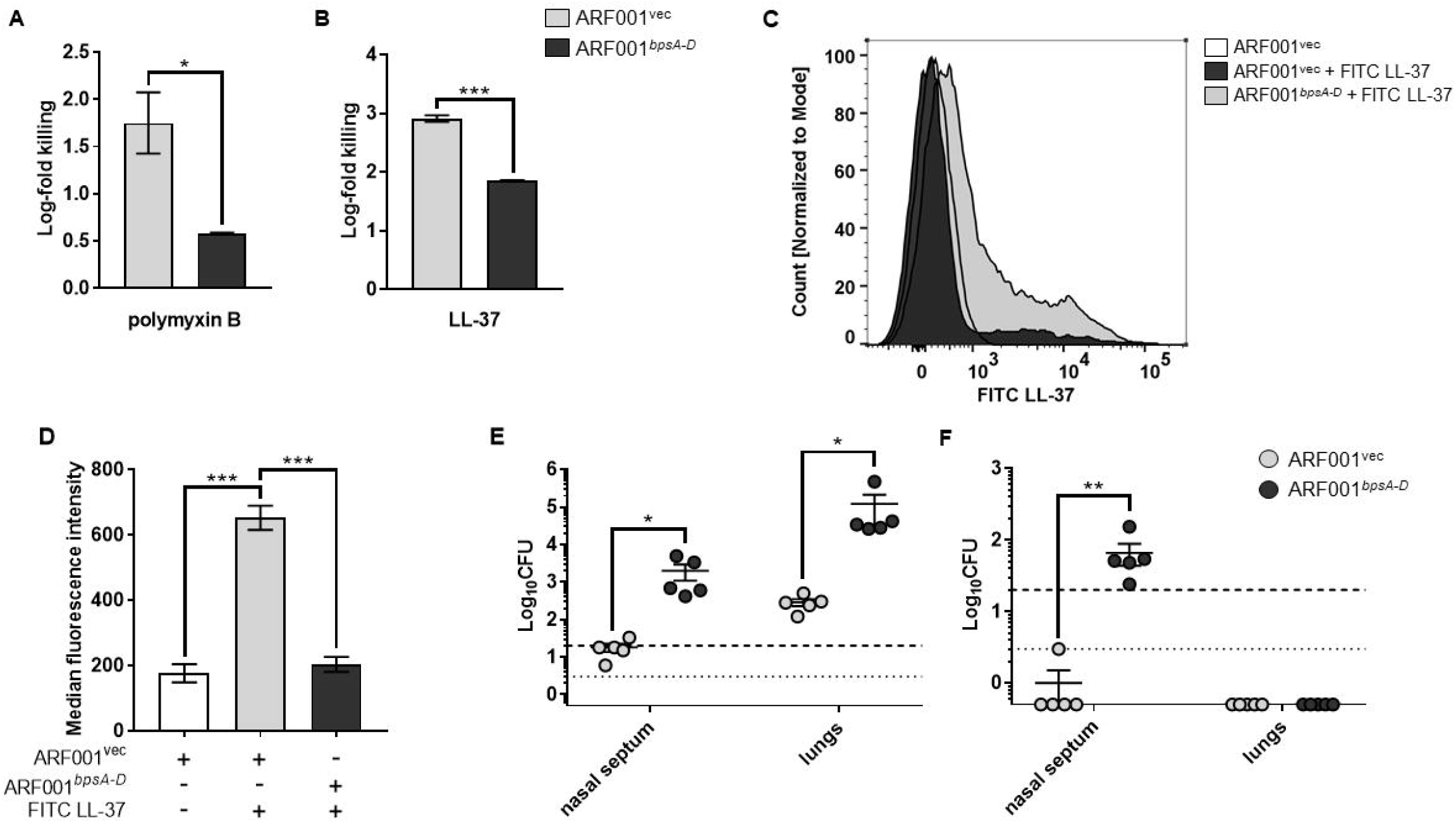
Bps enhances PmB and LL-37 resistance, inhibits AMP binding, and promotes respiratory tract survival when produced in *E. coli.* (a, b) Survival of ARF001^vec^ and ARF001^*bPsA-D*^ strains in the presence of 0.5 μg/ml polymyxin B (a) or 0.5 μg/ml of LL-37 (b). Bacteria were exposed to the indicated concentrations of AMPs for 2 hours in 10mM Na3PO4 buffer. Each data point represents the mean and s.e.m. of triplicates from one of two independent experiments. Statistical differences were assessed by unpaired two-tailed Student’s *t* test. *, p<0.05; ***, p<0.0005. (c, d) Binding of FITC LL-37 to the ARF001^vec^ or ARF001*^bpsA-D^* strains was measured by flow cytometry. Bacteria were fixed and exposed to FITC-labeled LL-37 and then the fluorescence intensity of FITC-labeled LL-37 bound to bacteria was measured by flow cytometry. (c) Bacterial counts for each strain were normalized to mode. (d) Median fluorescence intensities were quantified and assessed for statistical analysis. Data represent triplicates from one of two independent experiments. Statistical differences were assessed by one-way ANOVA. ***, p<0.0005. (e, f) Bacterial CFUs recovered from the nasal septum and lungs three days after intranasal challenge (e) or one day after aerosol challenge (f) with either the ARF001^vec^ or ARF001^*bpsA-D*^ strains. Bars indicate the mean and s.e.m. of groups of five mice each. Data are representative of one of two independent experiments with five mice each. Statistical differences were determined by unpaired two-tailed Student’s *t* test for each organ. *, p<0.05; **, p<0.005. Dotted line represents the lower limit of detection for nasal septum, and dashed line represents the lower limit of detection for lungs.

C57BL/6 mice were infected utilizing both intranasal and aerosol routes [32] with either the ARF001^vec^ or ARF001^*bpsA-D*^ strain and CFUs were enumerated from the nasal septum and lungs four days post-challenge. Compared to mice infected with the ARF001^vec^ strain, mice intranasally infected with the ARF001^*bpsA-D*^ strain had approximately 100- and 400-fold higher bacterial burden in the nasal septum and lungs, respectively (Fig. 6E). Upon infection of mice by the aerosol route, the ARF001^vec^ bacteria were recovered from the nasal septum of only one mouse at the lower limit of detection (3 CFUs) (Fig. 6F). No bacteria were recovered from the lungs of any of the mice infected by aerosol. This suggests that ARF001^vec^ strain is unable to survive in the mouse nose and lungs when infected by the aerosol route. In comparison, the ARF001^*bpAD*^ bacteria were recovered from the nasal septum when infected by the aerosol route (Fig. 6F). The observed decrease in colonization in the nasal septum and lungs after aerosol infection was not due to the inability of bacteria to reach these sites as enumeration of CFUs approximately 30 minutes after aerosol infection resulted in the recovery of both the ARF001^vec^ and ARF001^*bpsA-D*^ strains from the nasal septum and lungs (Fig. S3).

Evaluation and semi-quantitative lesion scoring performed on lungs from mice intranasally infected with ARF001^vec^ (Fig. 7A-D) strain revealed mild neutrophilic interstitial pneumonia, mild to moderate thickening of the pulmonary interstitium, and moderate numbers of neutrophils infiltrating the interstitium and bronchioles. In contrast, lungs from mice infected ARF001^*bpsA-D*^ (Fig. 7E-H) were characterized by a marked neutrophilic and macrophagic pneumonia with significant regional consolidation and thickening of the interstitium. In central areas of consolidation, very large numbers of neutrophils and macrophages obscured the distinction between alveoli and interstitium, and multifocally, alveolar walls were lined by hyperplastic type II pneumocytes, a common pulmonary response to injury. BALT was similarly expanded in this group of lungs adjacent to bronchioles. Degeneration and necrosis, edema, and hemorrhage were similar in all examined lung specimens. Total histopathology scores from mice infected with ARF001^*bpsA-D*^ were significantly higher than total scores from mice infected with ARF001^vec^ (Fig. 7I). Taken together, these results suggest that the production of Bps in *E. coli* is sufficient to impart the ability to colonize the mouse respiratory tract and induce pathology in the lungs.

**Figure 7.**
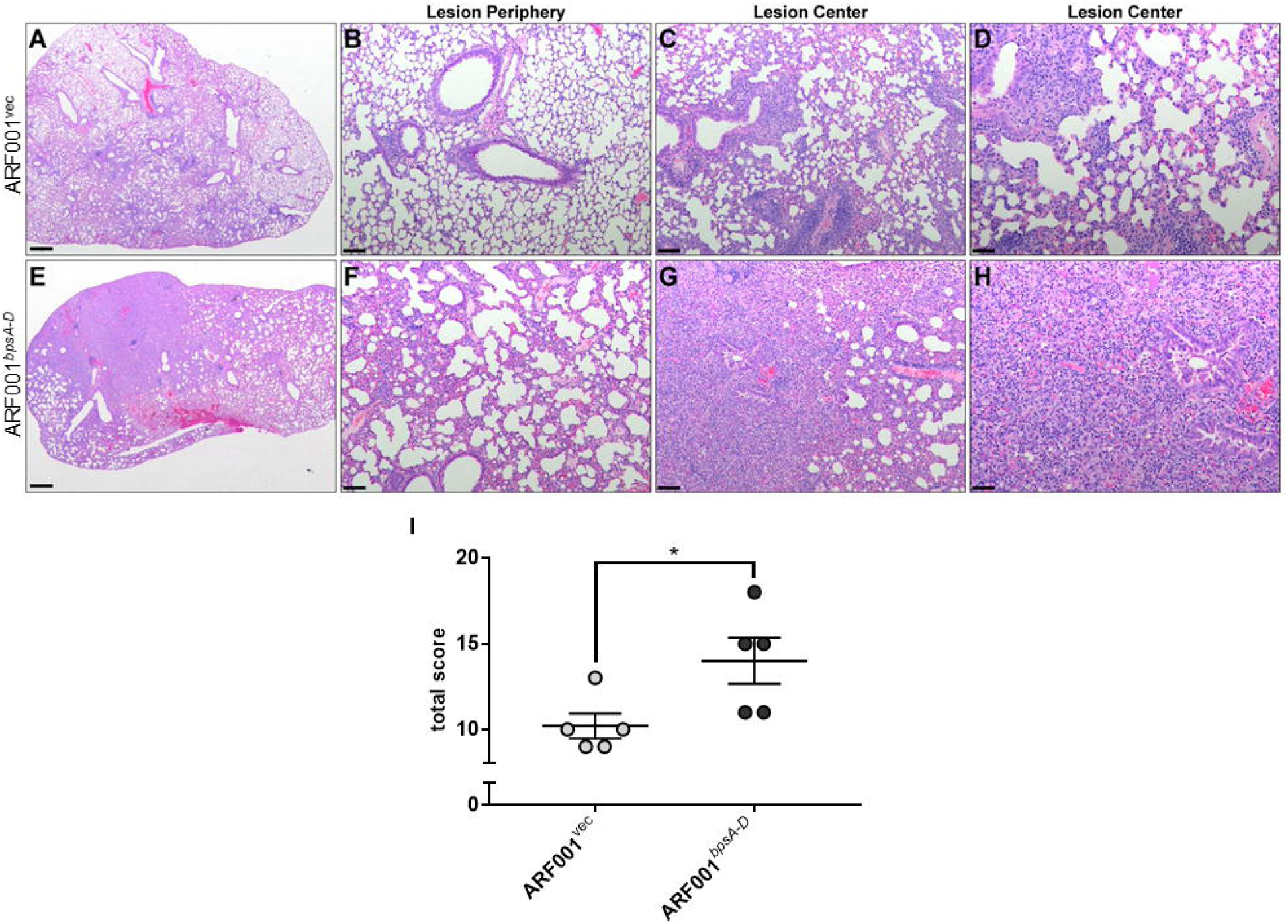
Light micrographs of mouse lungs after *E. coli* infection. Pulmonary infection with ARF001^*bpsA-D*^ results in severe pneumonia characterized by severe regional consolidation and interstitial pneumonia at the periphery of examined sections (e-h) compared to ARF001^vec^ (a-d). Lungs from mice infected with ARF001^*bpsA-D*^ have numerous infiltrating neutrophils mixed with mild amounts of edema and multifocal hemorrhage in the periphery of the tissue (f), while consolidation in the center of the lesions consists of very large numbers of neutrophils and macrophages, which obscure the distinction between alveoli and interstitium. Multifocally, there are areas of type II pneumocyte hyperplasia. Additionally, respiratory epithelium lining bronchioles is markedly hyperplastic to dysplastic or occasionally necrotic, with blebbing and sloughing of cells into the lumen and infiltration of neutrophils into the epithelium and lumen (g, h). In comparison, lungs from mice infected with ARF001^vec^ have only a few leukocytes in the interstitium peripherally (b), and lesion centers are characterized by small areas of inflammation and minimal consolidation around bronchioles (c, d). Inflammatory leukocytes are overall comparable between groups in that neutrophils are predominant and mixed with macrophages, however there are many more neutrophils in lungs infected with ARF001^*bpsA-D*^. Scale bar in a and e = 500 μm, 20x total magnification; scale bar in b, c, f, g = 100 μm, 100x total magnification; Scale bar in d and h = 50 μm, 200x total magnification. (i) Total histopathology scores of murine lungs infected with the indicated *E. coli* strains. Bars indicate the mean and s.e.m. of one representative of two independent experiments consisting of groups of five mice each. Statistical differences were assessed by unpaired two-tailed Student’s *t* test. *, p<0.05.

## Discussion

Successful prevention of microbial infection and disease necessitates a deeper understanding of the mechanisms by which microorganisms avoid host immunity. AMPs constitute a major component of the first line of immune defense in the mammalian respiratory tract. However, the respiratory pathogen *Bp* multiplies rapidly upon natural human infection and experimental infection of laboratory animals, and is thus capable of overcoming these defenses [33, 34]. In this manuscript, we investigated the role of the Bps polysaccharide in protecting against AMP-mediated killing. We demonstrated that compared to the WT strain, the isogenic *ΔbpsA-D* mutant strain was more sensitive to several structurally diverse human AMPs. Treatment of the WT strain by DspB, a glycoside hydrolase that specifically cleaves Bps, also enhanced AMP sensitivity.

We have discovered two different strategies by which Bps confers resistance to AMPs. First, we demonstrated that the *ΔbpsA-D* mutant strain bound higher amounts of LL-37 than the WT strain. This suggests that the presence of Bps on the cell surface inhibits AMP-bacteria interactions. For many Gramnegative bacteria, the LPS O-antigen plays a critical role in AMP resistance by limiting their interactions with the bacterial surface. However, *Bp* does not produce O-antigen [35]. We propose that in the absence of the O-antigen, Bps functions as a protective shield and blocks the bactericidal effects of AMPs on the bacterial cell.

Second, we report that *Bp* secretes Bps both in the medium during laboratory growth and in mouse lungs during infection. Incubation with both naturally secreted and purified Bps increased the resistance of bacterial cells to PmB and LL-37. Cell-free Bps bound to both PmB and LL-37. This suggests that cell-free Bps can sequester/trap AMPs by functioning as a sink and thereby neutralize their bactericidal activity. This property of Bps resembles that of decoy receptors found in mammalian systems, which bind specific growth factors or cytokines [36, 37]. Rather than signaling or activating the receptor complex this binding resulted in inhibition because the signaling molecules were trapped and rendered inactive. While many PNAG polysaccharides are secreted during laboratory growth [8, 38, 39], this is the first time a naturally secreted member of the PNAG polysaccharide family has been found to be secreted during infection and to mediate protection against a host immune component by functioning as a decoy.

PNAG/PGA polysaccharides contain variable amounts of de-*N*-acetylated glucosamine residues, rendering these polymers positively charged [8, 38, 39]. The deacetylation status of Bps is not known. Previously, the PIA polysaccharide of *Staphylococcus epidermidis* was shown to protect against killing by LL-37, and it was suggested that electrostatic repulsion is a likely mechanism of this protection [38]. However, this hypothesis has not been experimentally tested. Thus, it is noteworthy that cell-free Bps bound to positively charged AMPs and neutralized their killing activity. A previous study investigated the conformations of a series of linear and cyclic oligosaccharides related to PNAG. Linear oligosaccharides were not rigid and adopted several different conformations. The cyclic di-, tri, and tetra-saccharides adopted a symmetrical flattened ring conformation whereas the larger cyclic oligosaccharides were characterized by complicated shapes, resembling twisted rings [40]. We propose that the binding of AMPs to Bps and other PNAG polysaccharides is likely to be variable and may involve molecular forces other than ionic forces.

Previously when inoculated intranasally with single strains and compared to the WT strain, the *ΔbpsA-D* strain survived at considerably lower numbers in the mouse respiratory tract [6]. In the current report, we obtained similar results when mice were infected with single strains using the aerosol route. Further, we observed that when mice were infected by the aerosol route with a 1:1 mixture of the WT and *ΔbpsA-D* strains, the numbers of the mutant strain harvested from the nasal septum and lungs were similar to that of the WT strain. We also observed increased survival of the mutant strain when incubated with LL- 37 in the presence of the WT strain. We propose that both cell-associated and secreted Bps can bind and sequester AMPs present in the mouse respiratory tract and thereby enhance the survival of the mutant strain.

Production of Bps as a single *Bp* virulence factor in a laboratory-derived non-pathogenic *E. coli* strain conferred the ability to resist killing by PmB and LL-37. Like that observed in *Bp*, production of Bps inhibited AMP binding to *E. coli.* Production of Bps also increased the survival of *E. coli* in the mouse nose when mice were infected by the aerosol route, and in both the nose and lungs when infected by the intranasal route. Histological analyses of the lungs showed marked enhancement of pathology when mice were intranasally infected with the Bps-producing *E. coli* strain. We consider these findings to be quite striking, since continuous laboratory passage and culturing of *E. coli* has led to several genetic mutations, deletions, and loss of surface structures resulting in a non-pathogenic organism which is not suited to a life inside the host or outside a laboratory [41, 42]. Furthermore, *E. coli* is rarely considered to be a respiratory bacterium, as most pulmonary infections by *E. coli* are a result of dissemination from the gastrointestinal or urinary tract and from oropharyngeal aspirations [43].

In conclusion, Bps promotes resistance against AMP-mediated killing of *Bp* by functioning as a dual surface shield and decoy. The current findings further corroborate Bps as a principal determinant of *Bp* virulence and should serve as a model for investigating the pathogenic roles of other PNAG polysaccharides, which are conserved in many microbial species. Since Bps is not a component of current acellular pertussis vaccines, a vaccine containing Bps could control the resurgence of pertussis. Finally, Bps is a rare bacterial factor in that it confers virulence by itself in animal models of infection.

## Materials and Methods

### Bacterial growth conditions

Bacterial strains and plasmids used in this study are listed in Table S1. *Bp* strains were maintained on Bordet-Gengou (BG) agar supplemented with 10% defibrinated sheep blood (HemoStat, Laboratories, Dixon, CA) at 37°C for four days. For liquid cultures, *Bp* strains were grown in Stainer-Scholte (SS) broth supplemented with 0.1 mg/ml of heptakis (2,6-di-*O*-methyl-β-cyclodextrin, Sigma Aldrich, St. Louis, MO, USA) [20, 44, 45] at 37°C in a roller drum (80 rpm). *E. coli* strains were grown on either Luria-Bertani (LB) agar or in LB broth. As necessary, growth medium was supplemented with the appropriate antibiotics: chloramphenicol (Cm), 10 μg/ml; streptomycin (Sm), 100 μg/ml; and nalidixic acid (Nal), 20 μg/ml.

### Antimicrobial peptide killing assays

*Bp* and *E. coli* strains were grown to an OD_600_=1.0 in SS or LB broth, respectively. Bacterial cells were harvested by centrifugation (13,500 rpm, 5 min), washed twice with sterile 1X PBS, and resuspended in 10mM sodium phosphate buffer (pH 7.0). Dilutions of AMPs were prepared in 10mM sodium phosphate buffer, and bacterial cells were incubated with indicated concentrations of AMPs rotating at 37°C for 2 hours. CFUs were enumerated by spotting 10 μl of 10-fold serial dilutions onto BG agar plates with Sm (*Bp*) or LB agar plates (*E. coli)* with Cm. Log-fold killing was determined by subtracting the log10 CFU recovered after AMP treatment by the log10 CFUs recovered from only buffer added controls. Percent survival was determined by dividing the number of CFUs recovered after AMP treatment by the number of CFUs recovered from only buffer added controls.

For co-culture killing assays, WT and *ΔbpsA-D* strains were mixed in a ratio of 1:1 in 10mM sodium phosphate buffer (pH 7.0) and incubated with LL-37 at 37°C rotating for 2 hours. We recently discovered that while the WT strain is resistant to both streptomycin and nalidixic acid, absence of the *bpsA-D* locus results in sensitivity to nalidixic acid (Fig. S4). Therefore, we took advantage of the differential sensitivities of the two strains to nalidixic acid to track their survival in co-culture experiments. Bacterial suspensions were plated on both BG agar with Sm (to enumerate both WT and *ΔbpsA-D* strains) and BG agar with Nal (to enumerate only the WT strain). CFUs of *ΔbpsA-D* strain were calculated by subtracting the counts obtained on BG agar with Nal from the counts obtained on BG agar with Sm.

To determine the role of spent culture supernatant in AMP protection, bacteria grown to an OD_600_=1.0 in SS medium were pelleted by centrifugation (13,500 rpm, 5 min). After adjustment of pH to 7.4, the culture supernatants were filtered through a .22 μm filter (Millipore, catalog no. SCGP0052) and stored at −20°C. Bacteria were resuspended in either culture supernatants or SS broth. Bps prep or mock prep were purified as previously described [6]. AMP killing assays with *E. coli* strains were done under the same conditions as with *Bp*, and bacteria were enumerated on LB agar with Cm. For all AMP killing assays, at least two independent experiments were performed with n=3.

### Quantification of Bps by ELISA

*Bp* strains were grown to OD_600_=1.0), harvested by centrifugation (13,500 rpm, 15 min), washed with sterile 1X PBS twice and resuspended in sterile 1X PBS. To quantify cell-associated Bps, 1 x 10^8^ CFUs in 100 μl of PBS were incubated overnight at 4°C in 96-well plates (Corning, NY), washed three times with PBST (Tween 20) followed by blocking with 5% milk at 37°C for 1 hour. After washing three more times with PBST, the plates were incubated with wheat germ agglutinin (WGA) conjugated to horseradish peroxidase (HRP) (Biotium Inc. Hayward, CA, USA) at a dilution of 1:1000. After incubation at 37°C for 1 hour followed by five washes with PBST, 100 μL of 3,3’,5,5’- tetramethylbenzidine (TMB, Sigma Aldrich, St. Louis, MO, USA) were added to each well. The reaction was stopped using 100μl of 1M H2SO4.

To quantify secreted Bps, 1 ml of bacterial culture corresponding to OD_600_=1.0 was centrifuged (13,500 rpm, 15 min). Then, the supernatant was carefully aliquotted, filtered through a .22 μm filter (Millipore, catalog no. SCGP00525), and stored at −20°C for later use. 100 μl of supernatants were added to each well of 96-well plates. ELISA assays were performed as described above. Six independent experiments were performed with n=8.

To quantitate Bps from mouse lungs, the lung lysates from PBS-instilled and infected mice were centrifuged (5000 rpm, 5 min), the supernatant carefully aliquotted, filtered through .22 μm filter, and stored at −20°C for later use. 300 μl of supernatants were added to each well of 96-well plates and Bps was quantitated by ELISA as described above. Two independent experiments were performed with n=5. Two independent experiments were performed with n=5.

### Dispersin B treatment

Dispersin B (DspB) was purified as previously described [27]. Bacterial cells were harvested by centrifugation (13,500 rpm 5 min), washed twice with 1X PBS, and resuspended in DspB buffer (20mM Tris base, pH 8.0; 500mM NaCl). DspB treatments were carried out at indicated concentrations by incubating at 37°C rotating for 2 hours. The effect of DspB on cell-associated Bps was quantified by ELISA as described above. To perform AMP killing assays after treatment with DspB, bacteria were washed twice with 1X PBS, then resuspended in 10mM sodium phosphate buffer and incubated with indicated concentrations of AMPs. Two independent AMP killing assays were performed with n=3.

### Binding of LL-37 to bacteria by flow cytometry

Bacteria were harvested by centrifugation (13,500, 5 min), washed and fixed in 4% paraformaldehyde (PFA) with overnight rotation at 4°C. Cells were then washed with PBS, blocked with 1% BSA and probed with 1mM FITC-labelled LL-37 overnight rotating at 4°C. After two washes with PBS, approximately 10,000 events were collected per sample by using a Cytek Aurora Flow Cytometer (Cytek, Fremont, CA). FloJo software was used for data analysis. Two independent experiments were performed with n=3.

### Binding of AMPs to purified Bps

Serial dilutions of LL-37 were coated on high-binding 96-well plates (Corning, NY) and incubated at 4°C overnight. Next, plates were washed three times and probed with 50 μg/ml Bps and a mock preparation as described [6]. WGA conjugated to HRP (Biotium) was used to quantify the amount of Bps bound to LL-37 by ELISA as described above. Two independent experiments were performed with n=6.

For PmB binding, 10 μl of 1:2, 1:5 and 1:10 dilutions of 10 mg/ml stock of purified Bps preparations or mock preparations strain were spotted on a nitrocellulose membrane and allowed to dry overnight. The membrane was blocked in 5% milk for 30 min followed by incubation with 25 μg/ml of PmB for one hour. Next, mouse monoclonal anti-polymyxin B antibody (ab40014) (Abcam) was added at a dilution of 1:1000 for one hour followed by three washes with TBST. The HRP-conjugated secondary antibody was added at a dilution of 1:5000 for one hour, followed by three washes with TBST and detection by the ECL system.

### Animals

Housing, husbandry, and experiments with animals were carried out in accordance with the guidelines approved by the Institutional Animal Care and Use Committee of The Ohio State University. C57BL/6J mice (Jackson; male and female, 6 to 12 weeks old) were bred in-house. All experiments were reviewed and approved by The Ohio State University Institutional Animal Care and Use Committee (Protocol #2021A00000069).

### Mouse models of infection

Male and female C57BL/6J (Jackson) mice were used for all experiments. For all mouse experiments, two independent experiments were performed with groups of five mice.

For aerosol infection, mice were infected with 10^8^ CFU/ml of *Bp* WT and *ΔbpsA-D* strain, either as single strains or mixed in a 1:1 ratio, or the ARF001^vec^ or ARF001^*bpsA-D*^ as single strains, in an Allied Schuco^®^ S5000 Nebulizer for 30 minutes. A cohort of infected mice was sacrificed within 30 minutes after aerosol infection to determine the initial bacterial burden. After designated times post-challenge, the nasal septum and lungs were harvested, homogenized and *Bp* counts were enumerated on BG agar plates with Sm for single strain inoculum or by plating separately on BG agar with Sm and BG agar with Nal as described above. For mice infected with *E. coli* strains, CFUs were enumerated by plating on LB agar with Cm.

For intranasal infection with *E. coli,* 5×10^7^ CFUs in 50 μl (1X PBS) were intranasally administered to each mouse. After 3 days, the nasal septum and lungs were harvested from each mouse and bacterial counts were enumerated by plating LB agar with Cm.

A board-certified comparative pathologist (Dr. Corps) performed blinded semi-quantitative lesion scoring on n=5 lungs infected with *E. coli* strains and routinely stained with hematoxylin and eosin. Semi-quantitative lesion scores addressed the following parameters (Table S2) and were devised based on numerous previously published methodologies and in reflection of lesions present in the experimental cohort: degree of cellularity and consolidation of lung tissue as a percent of total tissue; thickness of alveolar walls; degeneration or necrosis in any portion of the examined lung; presence of edema; presence of hemorrhage; percent of examined alveolar and interstitial tissue infiltrated by neutrophils; percent of bronchioles infiltrated by neutrophils; percent of examined alveoli distant to the lesion center containing alveolar macrophages; and perivascular or peribronchiolar expansion of lymphoid populations +/- plasma cells (bronchioalveolar lymphoid tissue, BALT).

### Statistics

Statistical analyses of results were performed by unpaired two tailed t-test, one-way ANOVA, two-way ANOVA and Bonferroni posttest. All statistical analyses were performed using GraphPad Prism 7.05.

### Resource and Materials Availability

Further information and requests for resources, reagents, and protocols should be directed to and will be fulfilled by Lead Contact, Dr. Rajendar Deora. All unique reagents generated in this study are available from the Lead Contact with a completed Materials Transfer Agreement.

## Acknowledgements

RD and PD are supported by grants 1R21AI156732 and 1R01AI153829-01A1 from NIAID. This work was also supported in part by grants from the Canadian Institutes of Health Research (CIHR) to PLH (FDN154327). PLH is a recipient of a Tier I Canada Research Chair. Dr. Corps and the CPDISR are supported in part by grant P30 CA16058, National Cancer Institute, Bethesda, MD. ARF was supported in part by The Ohio State University fellowship program for Advancing Research in Infection and Immunity. We thank Drs. Daniel Wozniak and Lauren Bakaletz for critical reading of the manuscript, Dr. Mark Nitz for scientific discussions, and Dr. Landon Locke for the generous gift of FITC LL-37.

## Author Contributions

Conceptualization, Methodology, Validation, Formal Analysis, Visualization – ARF, JLGF, KSY, CFL, KNC, RD. Investigation – ARF, JLGF, KSY, CFL, HGC, MAV, DR, KNC. Writing – ARF, JLGF, KSY, KNC, PLH, RD. Resources, Supervision, Project Administration, Funding Acquisition – KNC, PLH, PD, RD.

## Declaration of Interests

The authors declare no competing interests.

---

**Fig. S1. Incubation with Dispersin B alone has no significant effect on the survival of *Bp*.** Survival of WT and *ΔbpsA-D* strains after incubation for 2 hours with Dispersin B at 37°C. Each data point represents the mean and s.e.m. of triplicates from one experiment and is representative of two independent experiments. Statistical differences were assessed by unpaired two-tailed Student’s *t* test.

**Fig. S2. Bacterial CFUs recovered from the nasal septum and lungs of mice 30 min after aerosol infection with *Bp.*** Bacterial CFUs recovered from the nasal septum and lungs of C57BL/6J mice 30 min after aerosol infection with co-culture of WT and *ΔbpsA-D* strain in a 1:1 ratio. Bars indicate the mean and s.e.m. of groups of five mice each. Data are representative of one of two independent experiments. Statistical differences were assessed by two-tailed Student’s *t* test for each organ. Dotted line represents the lower limit of detection for nasal septum, and dashed line represents the lower limit of detection for lungs.

**Fig. S3. Bacterial CFUs recovered from the nasal septum and lungs of mice 30 min after aerosol infection with *E. coli* strains.** Bacterial CFUs recovered from the nasal septum and lungs 30 min after aerosol challenge with either the ARF001^vec^ or ARF001^*bpsA-D*^ strains. Bars indicate the mean and s.e.m. of groups of five mice each. Data are representative of one of two independent experiments. Statistical differences were determined by unpaired two-tailed Student’s *t* test for each organ. Dotted line represents the lower limit of detection for nasal septum, and dashed line represents the lower limit of detection for lungs.

**Fig. S4. Bacterial growth on nalidixic acid.** WT and *ΔbpsA-D* strains were streaked on BG agar supplemented with 10% defibrinated sheep blood and 20 μg/ml nalidixic acid. Plates were incubated at 37°C for four days.

**Table S1. Strains and Plasmids.**

**Table S2. Histopathological Scoring Parameters.**

## Notes

### Competing Interest Statement

The authors have declared no competing interest.

