## Supplementary figures and images for "Bps polysaccharide of *Bordetella pertussis* resists antimicrobial peptides by functioning as a dual surface shield and decoy and converts *Escherichia coli* into a respiratory pathogen"

### Supplemental Figure 1

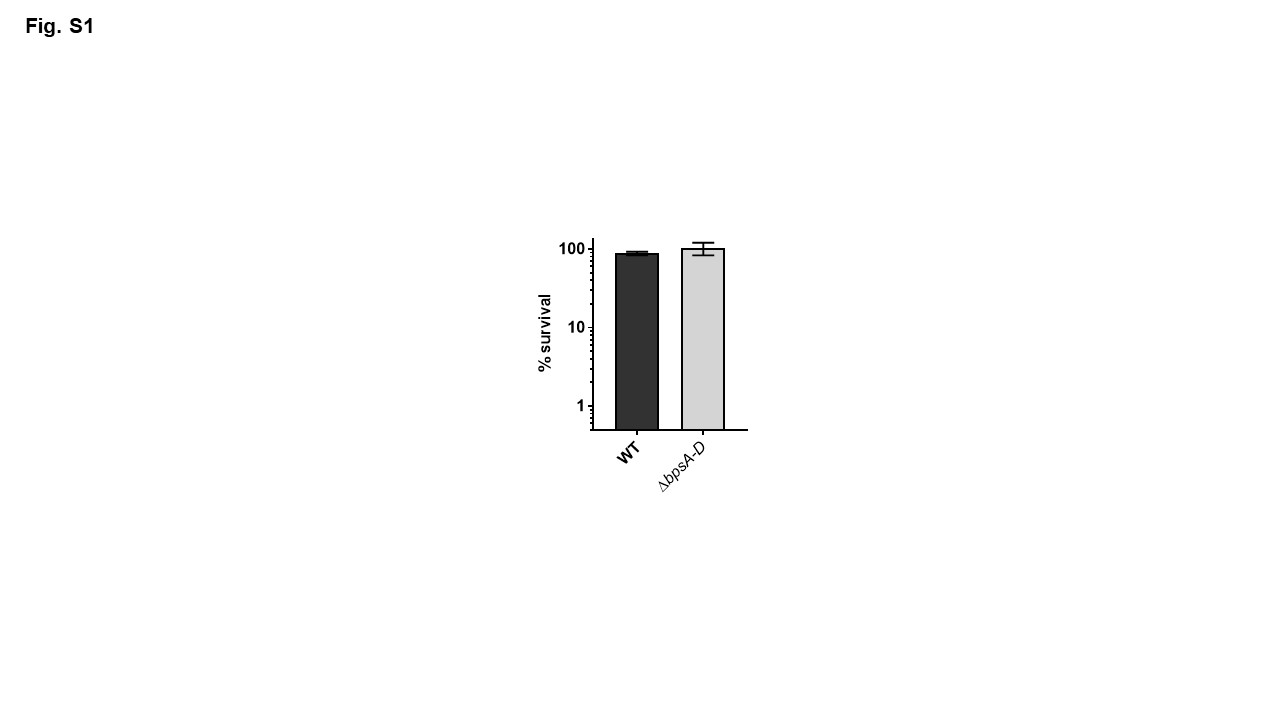

### Supplemental Figure 2

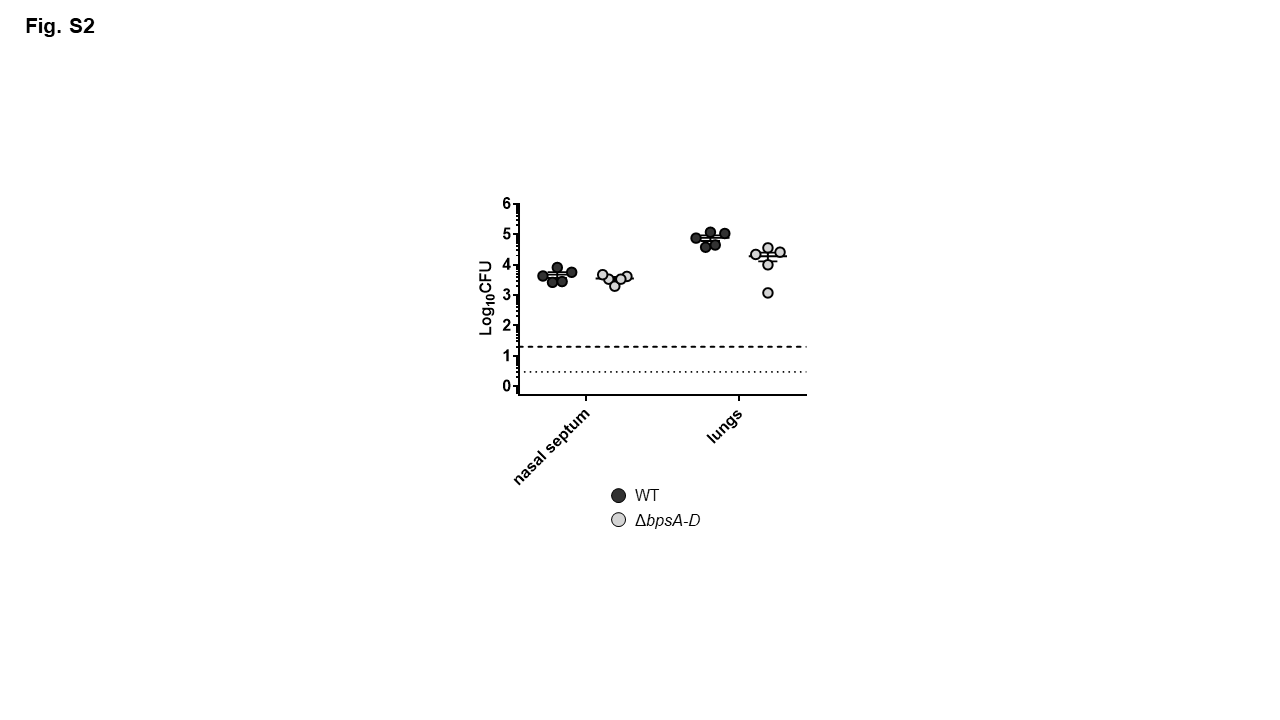

### Supplemental Figure 3

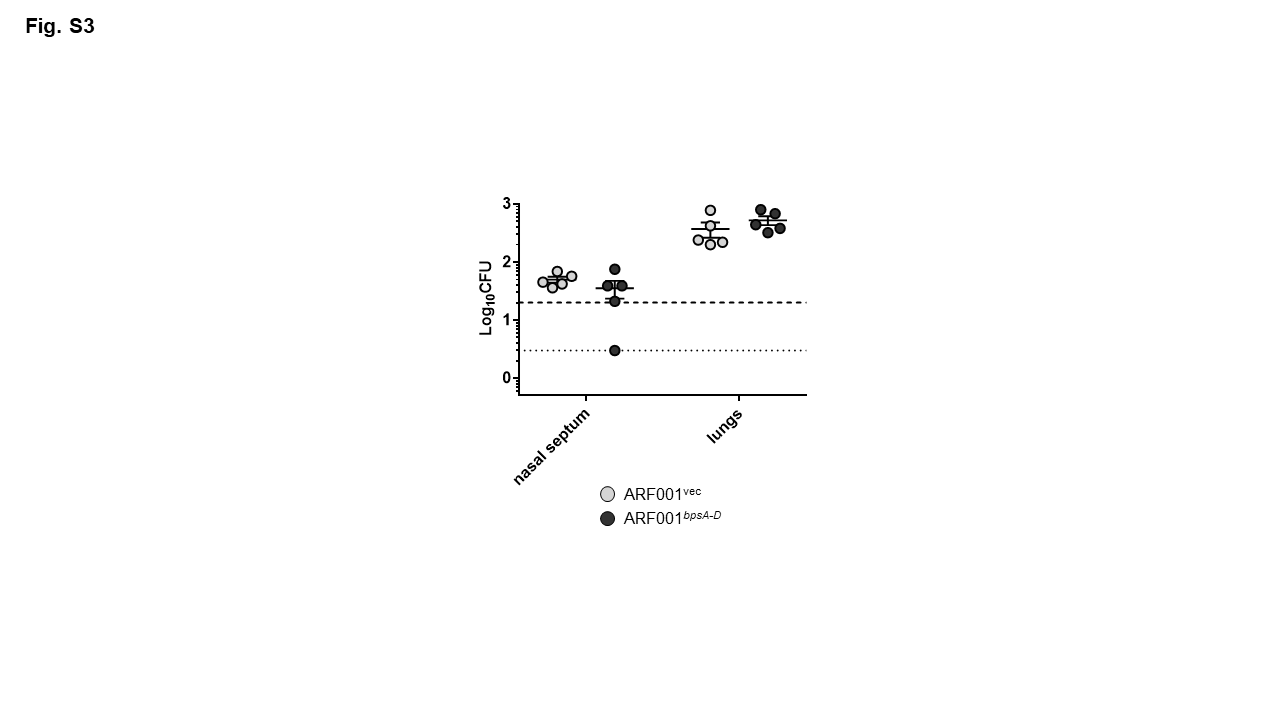

### Supplemental Figure 4

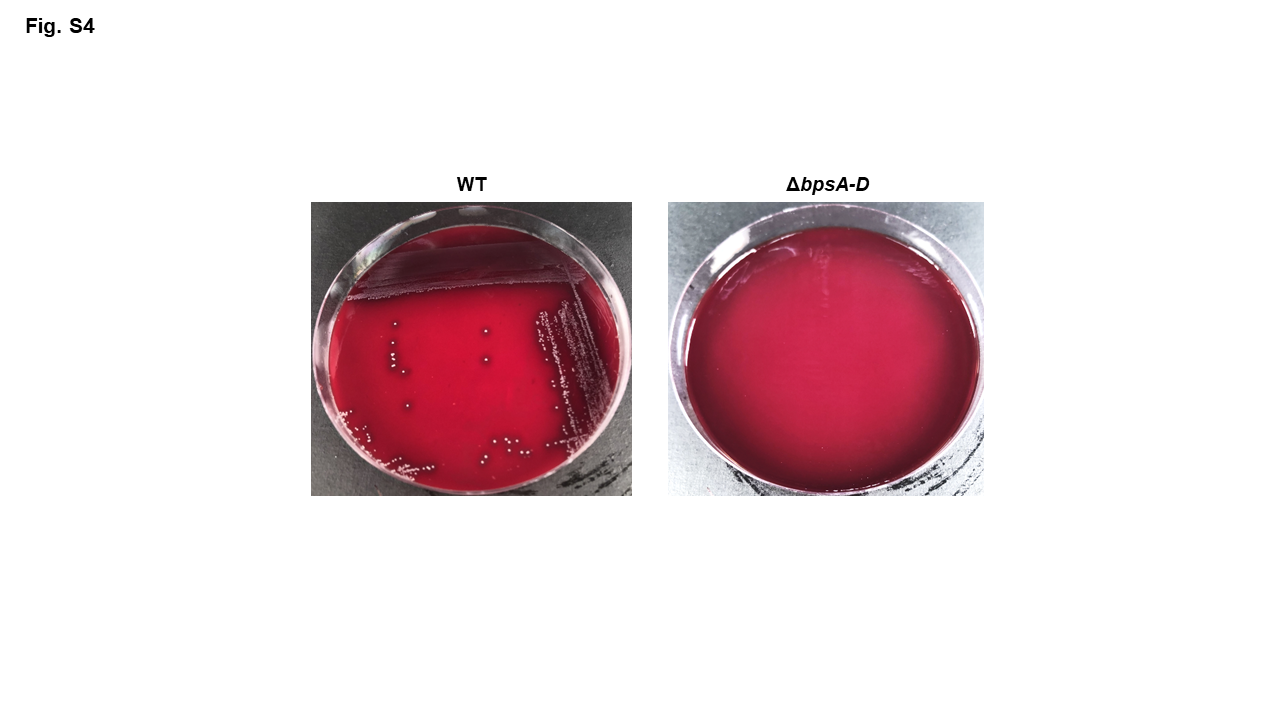

### Supplemental Table 1

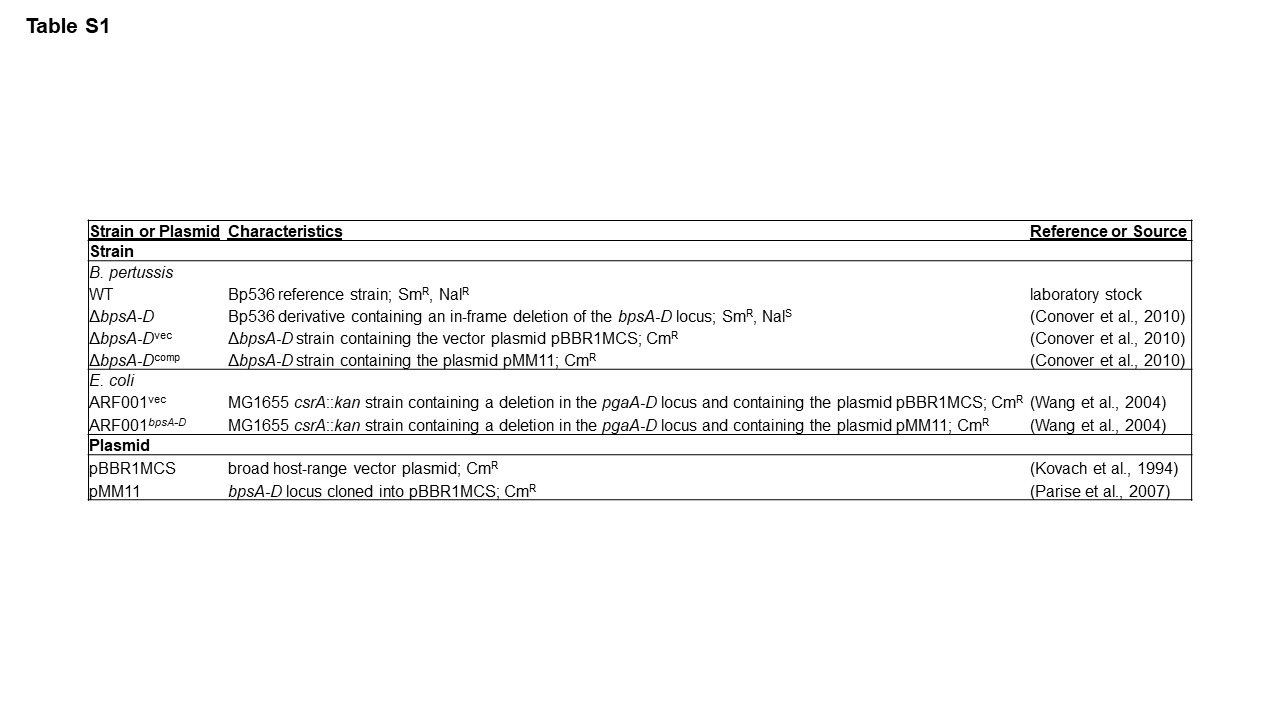

### Supplemental Table 2

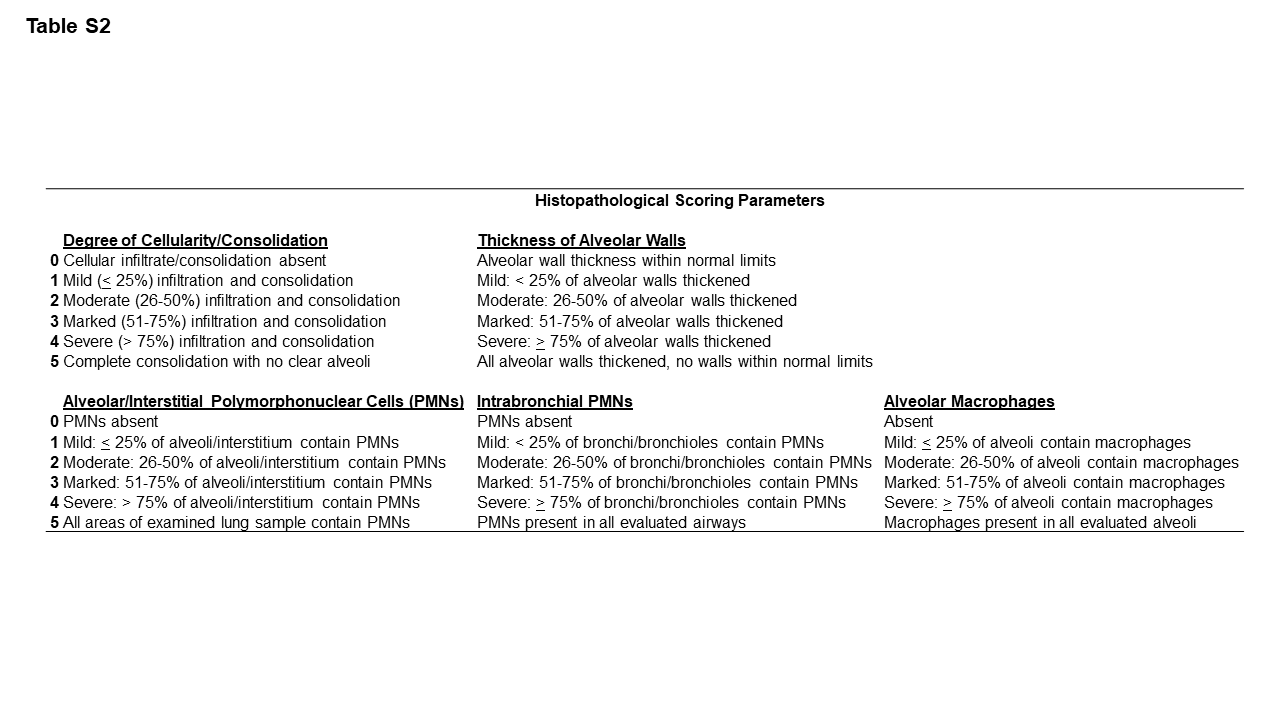
